# Deciphering gene regulatory programs in mouse embryonic skin through single-cell multiomics analysis

**DOI:** 10.1101/2024.10.11.617797

**Authors:** Qiuting Deng, Pengfei Cai, Yingjie Luo, Zhongjin Zhang, Wen Ma, Zijie Huang, Xiaoya Chen, Shijie Hao, Weiguang Ma, Jiangshan Xu, Mengnan Cheng, Xiumei Lin, Ru Zhou, Shanshan Duan, Junjie Chen, Ronghai Li, Xuyang Shi, Chang Liu, Peng Gao, Jianting Li, Jun Xie, Longqi Liu, Yue Yuan, Chuanyu Liu

**Author notes:** These authors contributed equally (Qiuting Deng, Pengfei Cai, Yingjie Luo, Zhongjin Zhang). Corresponding author: Longqi Liu (;) Yue Yuan; Chuanyu Liu.

## Abstract

**Background:** Cell type-specific transcriptional heterogeneity in embryonic mouse skin is well-documented, but few studies have investigated the regulatory mechanisms.

**Results:** Here, we present high throughput single-cell chromatin accessibility and transcriptome sequencing (HT-scCAT-seq), a method that simultaneously profiles transcriptome and chromatin accessibility. We utilized HT-scCAT-seq to dissect the gene regulatory mechanism governing epidermal stratification, periderm terminal differentiation, and fibroblast specification.

**Conclusions:** By linking chromatin accessibility to gene expression, we identified candidate *cis-*regulatory elements (cCREs) and target genes crucial for dermal and epidermal development. We described cells with similar gene expression profiles that exhibit distinct chromatin accessibility statuses during periderm terminal differentiation. Finally, we characterized the underlying lineage-determining transcription factors (TFs), and demonstrated that ALX4 and RUNX2 were candidate TF regulators of the dermal papilla lineage development through in silico perturbation analysis.

## Background

Embryonic mouse skin serves as an effective barrier, protecting the developing epidermis and dermis from environmental insults and amniotic fluid. Skin development starts post-gastrulation and experiences significant morphological changes before birth [1]. By embryonic day 9.5 (E9.5), a single layer of epidermal basal layer cells forms. Epidermis stratification and dermis maturation are primarily completed by E18.5, giving rise to multiple cell lineages [2, 3]. Current research mainly focuses on the epidermis of mouse skin [4, 5]. While the maturation of both dermis and epidermis involves diverse cell destinies, the intricate connections between cell lineage differentiation and gene regulatory networks (GRNs) in the epidermis and dermis are still need to be fully understood.

The transcriptome state indicates gene activation or repression, while epigenomic landscapes reveal potential cell type-specific CREs [6]. Advances in single-cell technologies have facilitated the exploration of factors determining cell identity within a tissue or organ. Single-cell RNA sequencing (scRNA-seq) is utilized to unveil the transcriptional dynamics during development of mouse skin and formation of the dermis [7–10]. The epidermis originates from surface ectoderm around E9.5 with a temporary cellular layer called periderm on the top of it. Then epidermis proceeds to stratify, and basal epidermal cells undergo differentiation. This periderm exists only for a brief period and sloughed off around E18.5 [11]. Additionally, around E14.5, the hair follicle placodes in the epidermal appendages emit signals to the dermal fibroblasts, leading to the formation of dermal condensates (DC), which are the precursor cells of the dermal papillae (DP). WNT and BMP signals within the dermal papillae trigger differentiation and maturation of hair follicles [12, 13]. While previous studies used scRNA-seq to reveal cellular heterogeneity in periderm and DP, but few is known about the epigenetic mechanism driving the specification of dermis [14, 15].

Lineage-priming of chromatin accessibility can indicate gene expression and predict lineage selection before commitment [16]. Parallel profiling transcriptome and chromatin accessibility within the same single cell can offer a global view of biological state rather than single-omics alone. However, current single-cell multiomics technologies are limited in throughput [17–19], sensitivity, and cost inefficient. Here, we present HT-scCAT-seq: **H**igh **T**hroughput **s**ingle-**c**ell **C**hromatin **A**ccessibility and **T**ranscriptome **seq**uencing, which enables simultaneous detection of chromatin accessibility and gene expression in thousands of single cells facilitated by a microfluidic device.

To systematically investigate cell-type specific gene regulatory programs in developing embryonic skin especially fibroblast within dermis, we applied HT-scCAT-seq to mouse embryonic skin. We profiled transcriptome and chromatin accessibility simultaneously in cells isolated from dorsal skin at E13.5, E14.5, E15.5, E16.5, and E18.5 stages. Our study reveals the dynamic heterogeneity within periderm and fibroblast populations during skin development. We identified crucial gene regulatory networks and characterized the underlying lineage-determining TFs in fibroblast lineage. Taken together, our work provides a comprehensive analysis of periderm and fibroblast population, and offering insights into epidermal and dermal development during embryogenesis.

## Results

### HT-scCAT-seq enables highly-accurate simultaneous chromatin accessibility and gene expression in single cells

In this study, we introduce HT-scCAT-seq, an enhancement of a previously published multi-omics approach, scCAT-seq [17]. HT-scCAT-seq enables robust detection of ATAC and RNA in fixed cells using microfluidics-based DNBelab C4 system [20]. The HT-scCAT-seq process involves five key steps: (I) nuclei are extracted and permeabilized using a mild lysis buffer and fixed with a low concentration of formaldehyde; (II) open chromatin regions are tagged by transposase Tn5; (III) mRNA transcripts are captured by a primer carrying poly (dT) and unique molecular identifiers (UMIs) during reverse-transcription; (IV) nuclei, beads, and nuclei lysis buffer is encapsulated into emulsion droplets facilitated by DNBelab C4 platform. The transposed chromatin fragments and RT products of the same single cell are barcoded by bead-linked single-strand oligonucleotides, and pre-amplified in droplets simultaneously; (V) the resulting product (RNA and ATAC partition) is separated by streptavidin beads followed by amplification with distinct primers to produce parallel libraries (**Fig. 1a and Additional file 1: Figure S1a**). The advantages of this approach include increased efficiency and specificity in capturing target molecules, as the biotin-streptavidin interaction enhances the recovery of RNA while minimizing non-specific binding. This can improve the overall sensitivity of our assays[16, 21]. However, we recognize potential drawbacks. The beads pull-down step may lead to the loss of some low-abundance RNA species, which could affect representation in the final libraries. For data processing, FASTQ libraries are de-multiplexed, then aligned to genomes. ATAC partition libraries perform beads calling and merging separately[22, 23]. Valid barcodes are transferred to RNA partition to perform bead merging, and RNA uses PISA [24] to generate gene matrices. Finally, the peak matrix is generated by peak calling. Then, peak and gene matrices are combined into an integrated matrix containing chromatin accessibility and gene expression information (**Fig. 1a**).

**Fig. 1.**
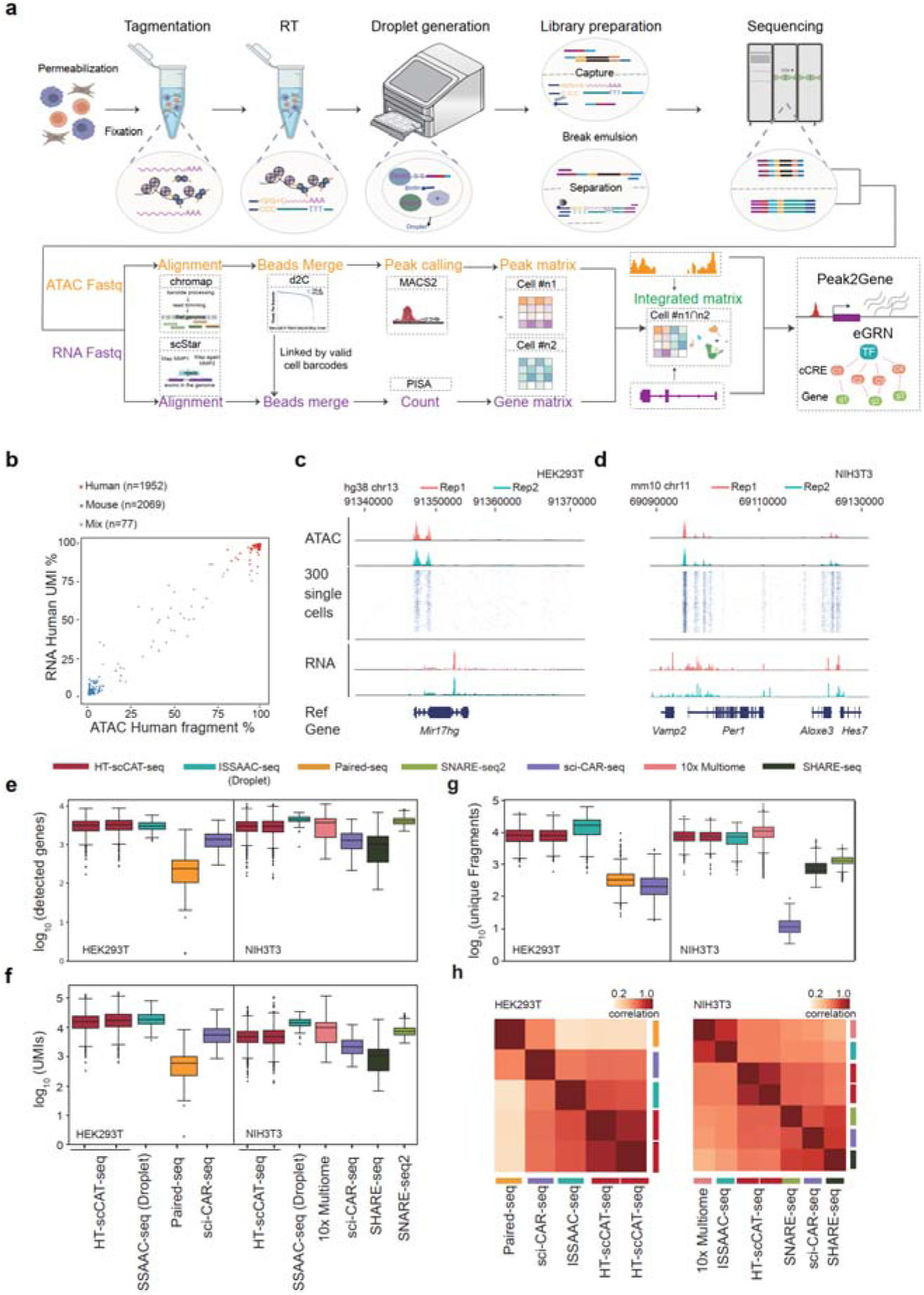
High-throughput single-cell chromatin accessibility and transcriptome sequencing based DNBelab C4 platform. **a** Scheme of the HT-scCAT-seq workflow and integrated data analysis. **b** Scatter plot of mixed-species experiment showing cells with both ATAC and RNA profiles obtained. Dots are colored based on species. **c**-**d** Track view displaying ATAC (top) and RNA (bottom) signals at a representative locus. c: *chr13: 91,340,000–91,370,000* for HEK293T cell. d: *chr11: 69,090,000–69,130,000* for NIH3T3 cells. The middle panel shows accessible chromatin fragments across top 300 selected single cells. **e-g** Box plots showing the distribution of detected gene number (**e**), UMIs (**f**) and FRiP (unique fragments in peaks, **g**) across different methods performed with HEK293T and NIH3T3 cells. **h** Heatmap showing the spearman correlation between transcriptome profiles generated by different approaches.

To evaluate its technical performance and data quality, we applied HT-scCAT-seq to an equal mixture of human HEK293T and mouse NIH3T3 cell lines. In this species-mixing experiment, human and mouse reads were well-separated with a collision rate (a portion of cells coincidently sharing same barcodes) of approximately 2% for both ATAC and RNA partitions (**Fig. 1b**). After filtering the doublets, 1,952 HEK293T cells and 2,069 NIH3T3 cells remained for analysis. We obtained a median of 3,679 RNA genes, and 11,556 unique ATAC fragments for each cell (**Additional file 2: Table S1)**. In HEK293T cells, reads from the ATAC and RNA library were highly enriched in the regions of *Mir17hg* gene loci. In NIH3T3 cells, reads from the ATAC and RNA library were highly enriched in the regions of *Vamp2*, *Per1*, and *Hes7* gene loci (**Fig. 1c, d**). To evaluate the ability of HT-scCAT-seq to capture chromatin accessibility, we compared the insert size distribution of unique ATAC fragments and mapped reads around transcription start sites (TSSs) from two biological replicates (**Additional file 1: Figure S1b-c**). As expected, reads from the ATAC partitions exhibited the expected periodical nucleosome pattern and a high enrichment around TSSs. For the ATAC and RNA partition, the ensemble signals revealed reproducibility (Pearson correlation coefficient >0.99) between two biological replicates (**Additional file 1: Figure S1d-e**).

When examining key metrics of ATAC libraries (unique fragments) and RNA libraries (UMIs and detected genes), we found that HT-scCAT-seq performed well in both modalities. This HT-scCAT-seq method yields data as high-quality as 11,396 ∼ 25,487 unique fragments (ATAC), 6,133 ∼ 8,430 UMIs, and 2,831 ∼ 3,816 detected expressed genes (RNA) per cell (**Additional file 2: Table S1**). We also benchmarked HT-scCAT-seq alongside other single-cell multiomics approaches [16, 21, 25–27]. The quality of data generated by HT-scCAT-seq is comparable to those by commercial 10x Multiome and ISSAAC-seq, which featured by detected UMI number, expressed gene number, and number of unique fragments (**Fig. 1e-g**). Global analysis reveals a high correlation between gene expression profiles from HT-scCAT-seq, 10x Multiome, and ISSAAC-seq (**Fig. 1h**). We also observed a high correlation between HT-scCAT-seq replicates (**Fig. 1h**), indicating stability of this approach. Taken together, these observations suggest that HT-scCAT-seq is sufficient to produce high-quality profiles of gene expression and chromatin accessibility in a reproducible and robust manner.

### Identification of mouse brain cell types from chromatin accessibility and gene expression profiles

To showcase HT-scCAT-seq’s effectiveness in identifying distinct cell types within complex tissues, we applied HT-scCAT-seq to adult mouse brain samples. After stringent quality filters, we generated single cell multimodal profiles of 14,828 high-quality mouse brain cells with a median of 6,991 UMIs and 5,471 unique fragments per cell (**Fig. 2a and Additional file 1: Figure S2a, 3a**). For the ATAC data, the nuclei that passed the quality control exhibited a median fragment count of 5,471, with 60.6% of fragments in peak (FRIP), and a TSS score of 6.64 (**Additional file 1: Figure S2b and Additional file 2: Table S2**). Regarding data of both modalities, HT-scCAT-seq performed comparably to the 10x Multiome and other published multi-omics assays (**Additional file 1: Figure S2a, b and Additional file 2: Table S2**).

**Fig. 2.**
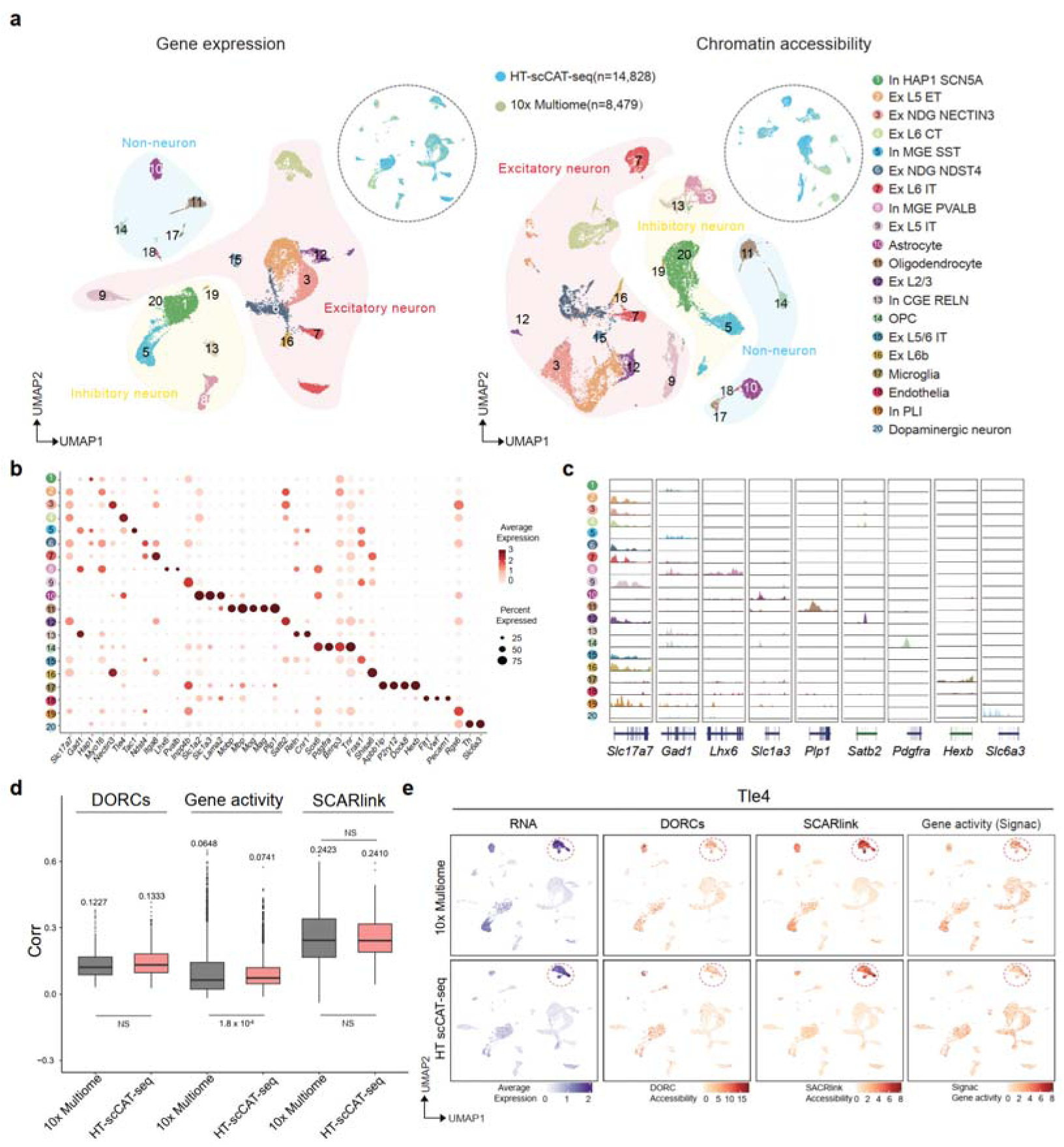
Evaluating the proficiency (efficacy) of HT-scCAT-seq in simultaneously capture ATAC and RNA profiles in the mouse brain. **a** UMAP visualization of RNA (left) and ATAC (right) profiles from 23,307 mouse brain cells, comprising both our data (n = 14,828) and 10x Multiome (n = 8,479). The cells are colored according to cell clusters (left) and approaches (top-right). Ex subtypes were identified by the neocortex areas (L2/3, L5, L5/L6 and L6). ET, extra-telencephalic; CT, corticothalamic; IT, intra-telencephalic; **b** Bubble plot showing expression level of neuronal/nonneuronal marker genes. **c** Aggregated scATAC tracks displaying signals in cell type-specific peaks. **d** Box plots displaying the Spearman correlation analysis between gene expression level and candidate CREs accessibility. DORCs: the Spearman correlation between the DORC matrix and gene expression values in the RNA dataset. Gene activity: the Spearman correlation between gene activity scores from ATAC dataset and gene expression values from the RNA dataset. SCARlink: the Spearman correlation between single-cell gene expression predicted by SCARlink, based on chromatin accessibility, and the actual gene expression values in the RNA dataset. The Spearman correlation was calculated by two-tailed Mann–Whitney U-tests. **e** UMAP colored by normalized gene expression, DORC score, SCARlink score and gene activity score of Ex_L6 CT marker *Tle4* (top: 10x Multiome, bottom: HT-scCAT-seq).

We further compared this dataset with a published mouse brain dataset generated by 10x Multiome [28] by integrating these two datasets. We projected cells based on RNA or ATAC profiles separately on two-dimensional UMAPs [29] and performed unsupervised clustering using RNA partition data. Then we transferred cell type labels which defined by expression of cell type specific marker genes [30], from RNA based clusters to corresponding ATAC profiles, and found that ATAC partition profiles generated by our approach or 10x Multiome recalled those clusters with minor differences (**Fig. 2a and Additional file 1: Figure S3a**). We revealed 9 excitatory neurons (Ex, *Nefh*, *Slc17a7*), 6 inhibitory neuron sub-types (In, *Nefh*, *Gad1*) and 5 non-neuron sub-types, including astrocyte (*Slc1a2*, *Slc1a3*), oligodendrocytes (*Plp1*), oligodendrocyte precursor cells (OPC, *Pdgfra*), microglia (*Hexb*), endothelia (*Pecam1*) (**Fig. 2b and Additional file 1: Figure S2c**). When looking into highest variable genes, aggregate ATAC signals within each sub-type display specific open chromatin peaks around the marker gene loci (**Fig. 2b, c and Additional file 1: Figure S3c**). Spearman correlation between each subtypes demonstrated a high level of congruence between the ATAC and RNA modalities (**Additional file 1: Figure S3d**). Cell type annotations were supported by gene expression and TF motif scores (**Additional file 1: Figure S3e, f**).

We introduced several computational algorithms to this mouse brain dataset to further compare the performance of HT-CAT-seq and 10x Multiome platforms. Three algorithms were employed in order to dissect gene expression – active *cis* regulatory element linkage (see Methods): (I) high density domains of regulatory chromatin (DORCs) score matrix was obtained by FigR [31], (II) gene activity was calculated with default parameter using GeneActivity function in Signac [32], with genes more than four associated peaks; (III) Single-cell ATAC + RNA linking (SCARlink) [33] was employed to predict single-cell gene expression using regularized Poisson regression based on single-cell chromatin accessibility data (see Methods). Comparable predictions scores of HT-CAT-seq and 10x Multiome dataset can be found when using either of these three algorithms (**Fig. 2d**). In our HT-CAT-seq data, predictions generated using gene activity score exhibited a significantly higher correlation with gene expression compared to the correlation calculated from 10x Multiome data (median corr = 0.0648 on 10x Multiome, median corr = 0.0741 on HT-CAT-seq). Furthermore, the correlation between DORC and SCARlink model predictions and gene expression from our data showed no significant difference compared to the 10x Multiome data (**Fig. 2d**). Conclusively, HT-scCAT-seq proves to be comparable to the 10x Multiome in its ability to predict gene expression based on multi-omic single-cell ATAC and RNA. Next, we examined the genes predicted by the FigR and SCARlink models and identified 114 genes shared by both models. We observed that Ex_L6 CT marker *Tle4* was identified more specifically in our data (**Fig. 2e**). As glutamatergic pyramidal neurons, Ex_L6 CT can elicit action potentials in layer 5a neurons while suppressing L4 neurons upon activation [34, 35].

### Single-cell multiomics reveals regulatory dynamics during mouse embryo skin development

To comprehensively understand the underlying regulatory mechanism driving the specification of multiple cell types during skin formation, we employed HT-scCAT-seq to embryonic mouse dorsal skin samples from E13.5 to E18.5 stages (**Fig. 3a**). After stringent quality control, cells from all five stages were analyzed together. Jointly analysis on both data modalities yielded a total of 64,408 cells with total 28,250 expressed genes and 175,103 accessible peaks (**Additional file 1: Figure S4a**).

**Fig. 3.**
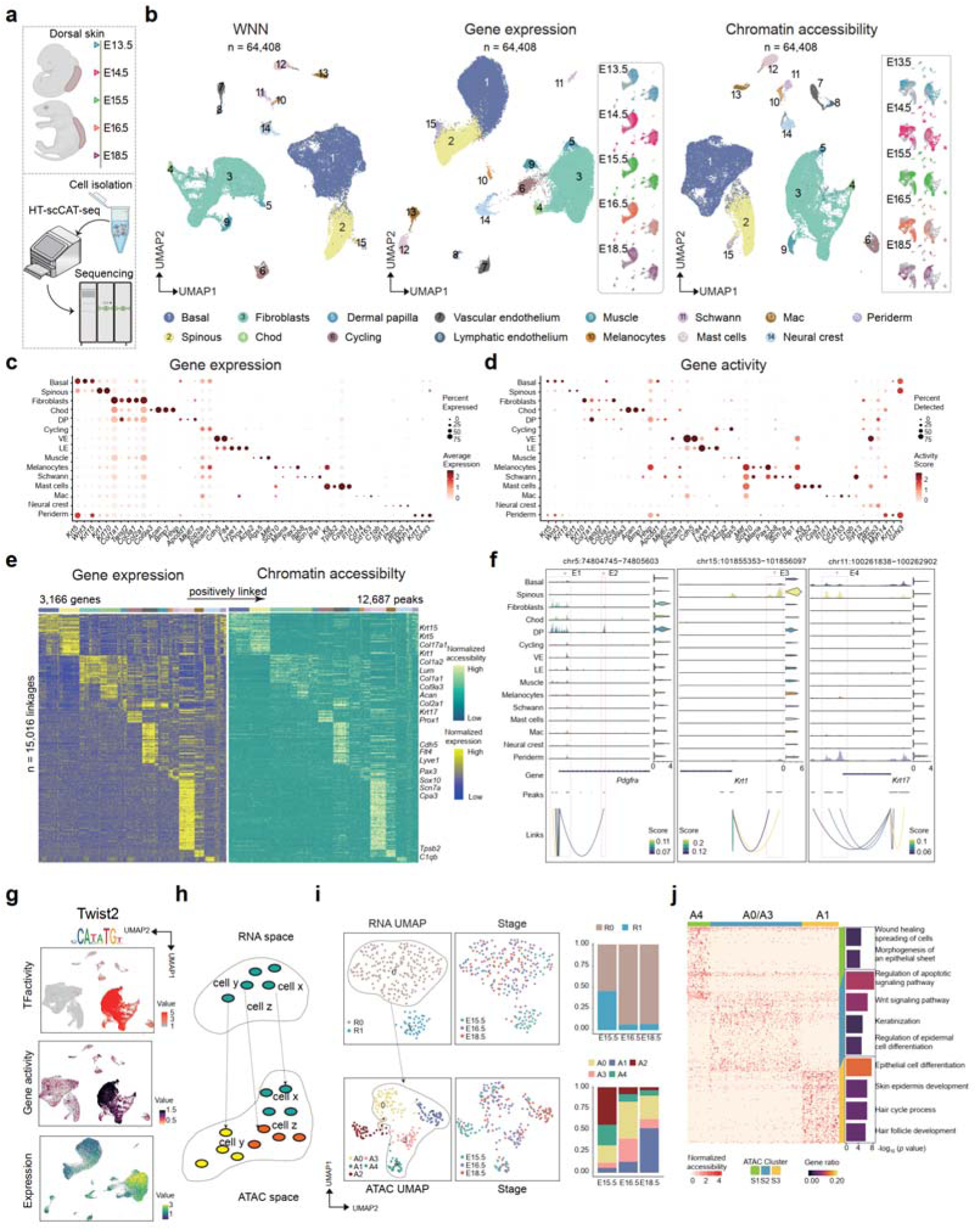
Single-cell multiomics assays uncovered dynamic features and heterogeneity of developing mouse skin. **a** Scheme of sample preparation. **b** UMAP visualization of WNN (left), RNA (middle) and ATAC (right) partition profiles, colored clusters. The dashed circle shows the same UMAP colored by developmental stage. **c** Bubble plot showing gene expression of selected marker genes for each RNA cluster. Color bar: relative expression levels across all clusters, bubble size: percentage of cells within each cluster that express the gene. **d** Bubble plot showing gene activity scores for the makers in (**c**). **e** Heatmap showing gene expression (left) and chromatin accessibility (right) for 15,016 significantly linked CRE-gene pairs. Each row represents a linked gene and a pair of CREs. Bar on the top represents the cell types involved in skin development. **f** Track view of aggregated ATAC signal around the *Pdgfra*, *Krt1*, *Krt17* locus. Peaks and peak-to-gene linkages are shown below the tracks. Right, violin plot shows the integrated expression levels of *Pdgfra*, *Krt1*, *Krt17* for each cell type. Red vertical bars highlight selected peaks linked to *Pdgfra*, *Krt1*, *Krt17* expression. **g** Feature plots showing TF activity score (top), Gene activity Score (middle)) and gene expression level (bottom) for *Twist2*. **h** Schematic of the conceptual workflow illustrating one state in RNA cluster corresponding to three states in ATAC cluster. **i** UMAP visualization of RNA (top) and ATAC (bottom) data from 591 periderm cells. Middle, nuclei are colored by the cluster (left) and developmental stage. Right, the distribution of periderm cells from E15.5 to E18.5 (right). **j** Heatmap showing normalized ATAC signals of the top 500 cluster specific peak peaks across three ATAC clusters. Right, representative enriched GO terms within each cluster.

We performed dimensional reduction of the resulting profiles using Seurat [36] and Signac [32], respectively. Biological replicates from the same developmental stage showed strong overlap in UMAP embedding (**Additional file 1: Figure S4b**). We performed cell type annotation using RNA partition datasets and transferred cell type labels to the corresponding ATAC clusters. Fifteen majors cell types were identified across nineteen distinct clusters (**Fig. 3b and Additional file 1: Figure S4c**). All expected embryonic skin cell types [7, 37] were verified by expression of canonical marker genes, including basal cell (*Krt5* and *Krt15*), spinous cell (*Krt1* and *Krt10*), periderm (*Grhl3* and *Myh14*), fibroblast (*Col1a1* and *Twist2*), dermal papilla (*Hhip* and *Bmp7*), vascular endothelium (VE, *Cdh5* and *Vwf*), lymphatic endothelium (LE, *Flt4* and *Acta2*), melanocytes(*Sox10*), neural crest (*Pcbp3*), schwann cell (*Itgb8*), muscle cells (*Rgs5*), mast cell (*Kit* and *Il1rl1*) and macrophages (*Cd163*) (**Fig. 3c, d**). Cells from different stages were well integrated, but their relative contributions to each cell type varied in agreement with temporal development of mouse skin [7, 15] (**Additional file 1: Figure S4d**). We calculated gene activity score [32] by summing up number of unique chromatin fragments intersecting gene body and promoter regions (**Fig. 3d**, see Methods). The above-mentioned RNA makers, also appeared similar patterns of chromatin accessibility in corresponding ATAC clusters, indicating strong congruence between these two modalities (**Fig. 3c, d**).

Next, we aimed to use multi-omics data to identify cell type-specific CREs and their target genes by comparing gene expression with chromatin accessibility across all cells in the dataset. We first identified specific gene expression for these cell types. Differential gene expression (DEG) analysis between subtypes revealed 5,299 DEGs (adjusted *p* value < 0.05 and log2(fold change) > 0.1). Feature linkages is characterized by a significant correlation between the accessibility of ATAC peaks and gene expression [38]. To identify regulatory elements correlated to each DEG, we performed a peak-gene linkage analysis by calculating the Pearson correlation coefficient (PCC) between gene expression and chromatin accessibility of peaks within 500 kb of each TSS [16, 39]. Positively correlated peak-gene pairs were identified as potential enhancer-gene interactions [40]. This analysis yielded a total of 15,016 peak-gene links, including 12,687 regulatory elements significantly linked to the 3,166 DEGs (**Fig. 3e and Additional file 3: Table S3**, correlation > 0, adjusted *p* value < 0.05). Each DEG was linked to a median of three peaks (min = 1, max = 39, mean = 4.743).

Then, we investigated whether candidate CREs could mediate the expression of differentially expressed genes. Our analysis revealed that some linkages with differentially expressed genes were uniquely identified in embryonic skin lineage cell types. For instance, the locus at chr5: 74,804,745−74,805,603 which mapped to the promoter region of *Pdgfra*, showed the most significant accessibility in dermal-derived cells (**Fig. 3f**). Compared to other cell types, dermal-derived cells (fibroblasts and DP) exhibited highest expression level of *Pdgfra*. Besides, we observed that a peak (E2, chr5:75,174,536-75,175,336) which linked with *Pdgfra* expression showed strong accessibility signal only in DP (**Fig. 3f)**. Together, we identify the locus at chr5: 74,804,745−74,805,603 as a candidate CRE for dermal-derived cells (fibroblasts and DP), capable of upregulating the expression of *Pdgfra* (**Additional file 1: Figure S4f**). Additionally, peak-gene links at *Krt17* (chr11:100,261,838−100,262,902) and *Krt1* (chr15:101,855,353−101,856,097) showed significant enrichment within periderm and spinous cells (**Fig. 3f and Additional file 1: Figure S4g, h**). *Krt17* is strongly expressed in periderm, whereas *Krt1* expressed in spinous cells. *Krt1* has been identified as a target of Notch signaling pathway in the epidermis and serves as a driver gene for the differentiation of basal cells into spinous cells [41–43].

To investigated the TFs potentially driving the regulatory programs in each embryonic skin cell type, we analyzed the enriched TF binding motifs that were present within these linked peaks using chromVAR (see Methods) [44]. The criteria for defining these peak gene-TF pair included a significant correlation between peaks and genes, high TF expression level, and enriched TF binding motif in accessible ATAC peaks [45]. This analysis resulted in the identification of 75 putative markers across 15 major cell types (adjusted RNA *p* value < 0.05 and adjusted motif *p* value < 0.05, **Additional file 4: Table S4**). For instance, motif enrichment analysis indicated a strong enrichment of TWIST2 binding motif in dermal-derived cells. We further demonstrated concordance by examining the RNA expression, gene activity score and TF activity of *Twist2* (**Fig. 3g**). *Twist2* serves as a marker of embryonic upper dermal fibroblasts [46]. *Twist2* Knockout in postnatal mice results in pronounced skin abnormalities including skin atrophy and fat deficiency, ultimately leading to death from cachexia within the first two weeks of life [47, 48].

To systematically investigate the gene regulatory program of lineage commitment during periderm development, we focused on the correlation between chromatin accessibility and gene expression in periderm cells. In our datasets, we examined periderm cells from five time points and classified them into early (E13.5 to E14.5) and late states (E15.5 to E18.5). Early periderm subcluster expressed *Tgfb2*, *Cldn23* and *Myh14*, while late periderm subcluster expressed markers of terminal differentiation such as *Bcl11b* (**Additional file 1: Figure S4i**). From E13.5 to E18.5, the cell from periderm cluster were consistently present in RNA UMAP, while in the ATAC cluster, they started to disappear from E15.5 (**Fig. 3b**). We conducted a focused analysis extracting 245 periderm cells from E15.5 to E18.5 and performed re-clustering. Two subclusters based on RNA profiles were identified (RNA clusters R0, R1), while five subtypes based on ATAC profiles were identified (ATAC clusters A0, A1, A2, A3, A4) (**Fig. 3h, i**). It suggests that chromatin accessibility may some part desynchronized with gene expression. (**Fig. 3j and Additional file 1: Figure S4j**). The heterogeneity was visible by clustering at the chromatin, showing five main chromatin states. To further investigate the epigenetic differences within the ATAC cluster, we conducted a focused analysis of cell from A0, A1, A3, A4. We identified three distinct chromatin accessibility statuses: S1, S2 and S3. The top 2,000 cluster-specific peaks, sorted by fold enrichment were taken for further analysis. Biological functions for peak associated genes of each status were annotated using genomic regions enrichment of annotations tool (GREAT) [49]. We noted that S1 status was enriched with functions such as wound healing, spreading of cells and epithelial sheet morphogenesis. S2 status was enriched with functions related to the regulation of apoptotic signaling pathway and keratinization, while S3 status was enriched with functions such as hair cycle process and hair follicle development (**Fig. 3j**). These results demonstrated that cells with similar gene expression profiles exhibit distinct chromatin accessibility status.

### Fibroblast heterogeneity and construction of GRNs governing fibroblast development

Dorsal dermis is a connective tissue derived from somatic mesoderm [50]. Dermis is embedded in an extracellular matrix (ECM) composed of collagen and elastic fibers. Among the developing dermis, fibroblasts are the major cell type that are considered an important cell lineage for skin development [51, 52]. While prior single cell studies have demonstrated dynamics of gene expression in fibroblasts, they provided limited insights on changes of chromatin state. We sought to utilize single-cell multiomics approach to identify fibroblast specific CREs and their target genes.

We first performed gene ontology (GO) analysis on DEGs of fibroblast-derived cells from E13.5 to E18.5, categorizing the fibroblasts into early (E13.5 to E14.5) and late (E15.5 to E18.5) stages (**Additional file 5: Table S5, Additional file 6: Table S6**). GO analysis reveals that early-stage fibroblast-derived cells were involved in regulation of the WNT signaling pathway, embryonic organ development, and connective tissue development [53]. These pluripotent functionalities may reflect early commitments towards future fibroblast fates (**Fig. 4a, Additional file 1: Figure S5a, Additional file 5: Table S5, Additional file 6: Table S6**). In the late-stage fibroblast-derived cells, the enriched GO terms included cellular response to lid, collagen fibril organization, and laminin interactions. These more specialized functionalities may indicate events related to fibroblast specialization [54] (**Fig. 4a and Additional file 1: Figure S5a**).

**Fig. 4.**
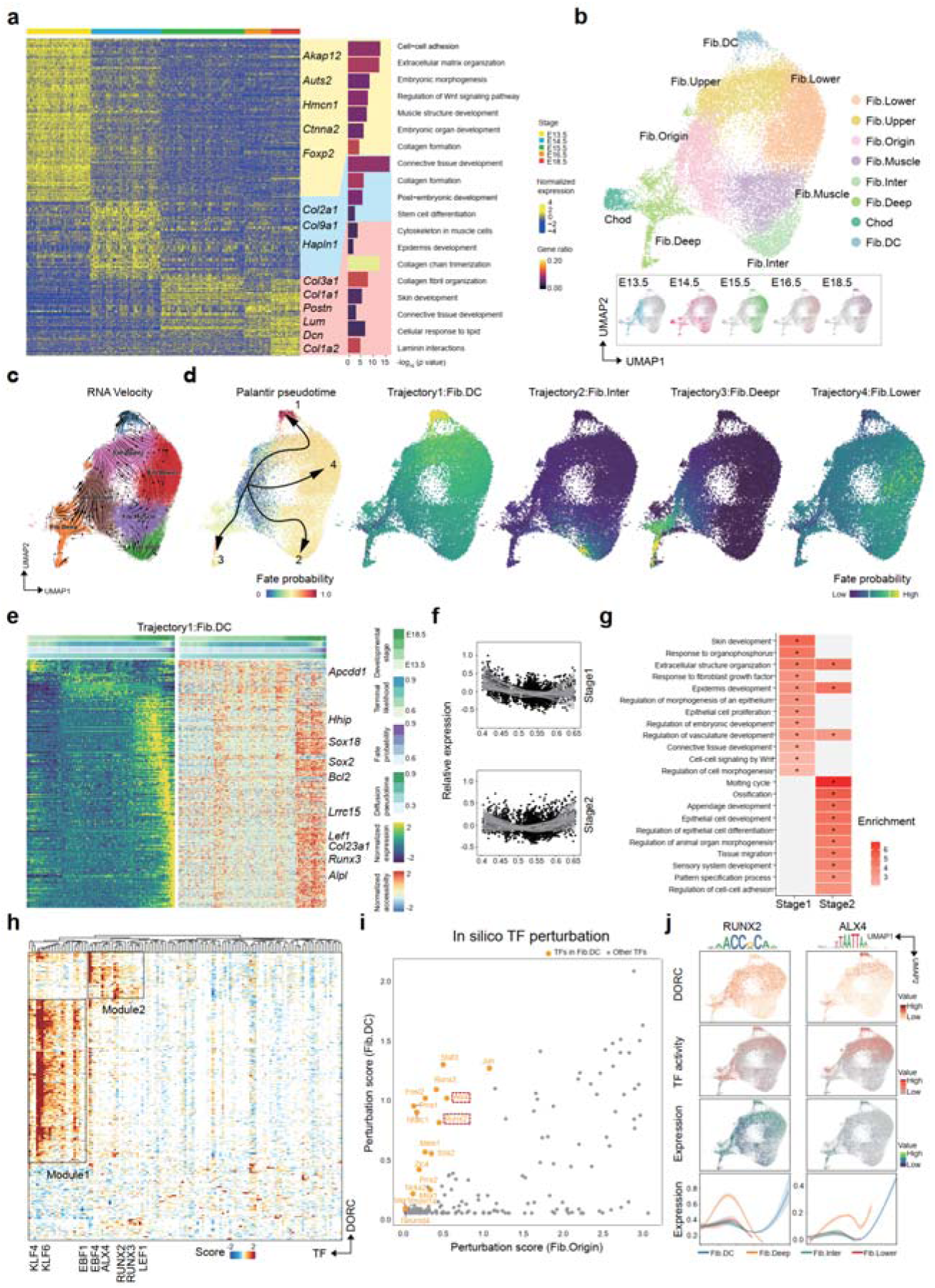
Reconstruction of fibroblast trajectories and identification of specific TF regulatory network. **a** Heatmap showing expression of DEGs of each developmental stage (adjusted *p* value < 0.05 and an average log_2_ fold change (log_2_FC) > 0.1), alongside a list of some well-studied markers. Histogram on the right shows enriched GO terms (molecular function) for each developmental stage. *p* value obtained from the hypergeometric test are shown, color scale indicates gene ratio. **b** UMAP visualizations of fibroblast subclusters using RNA profiles. Points are colored by cell types (top) or developmental stage (bottom). **c** RNA velocity of fibroblasts subtypes, projected onto the UMAP using scVelo. Colors indicate different cell types. **d** UMAP displaying cell fate bifurcation of fibroblast subtypes using CellRank. Four trajectories are shown: Fib.DC, Fib.Deep, Fib.Inter and Fib.Lower. Cells are colored according to fate probabilities by CellRank. **e** Heatmap showing gene expression (right) and chromatin accessibility (left) of 495 significantly linked gene-CRE pairs along the Fib.DC trajectory. Each row represents a pair of gene and a linked CRE. Columns are sorted by diffusion pseudotime. f. Scatter plots showing the gene expression values of driver genes·(y-axis) plotted against the pseudotime of the FibRC trajectory (x-axis). The fitted smooth line indicates a 95% confidence interval. **g** Enriched GO terms for driver gene sets upregulated at stage1 and stage2 of the Fib.DC trajectory. Dot size denotes the enrichment ratio、 and color reflects the level of significance. **h** Heatmap shows TF-DORC regulation scores derived from Fib.DC trajectory data using FigR (n= 125 TFs, n =395 DORCs), with DORC represented by row and candidate TF regulators by columns. **i** Scatter plot shows knockout (KO) simulation result of TFs in the Fib.Origin (x axis) and Fib.DC lineage (y axis). **j** UMAP colored by normalized DORC score, TF activity score, and gene expression level of underlying TFs in Fib.DC lineage (left: RUNX2, right: ALX4). The line plot (bottom) shows smoothed gene expression trends in Fib.DC trajectory.

To elucidate the heterogeneity within fibroblast cells, we extracted fibroblast profiles and re-clustered them into eight subtypes defined as previous reported [7] (**Fig. 4b**). We observed a continuous progression from E13.5 to E18.5 when fibroblasts are projected into low-dimensional subspaces (**Fig. 4b**). Cells from E13.5 were located near the center of the low-dimensional manifold, whereas cells from later stages were positioned towards the periphery, likely indicating more differentiated cell states (**Fig. 4b**). We observed distinct gene dynamics across eight fibroblast subclusters within our dataset (**Fig. 4b-d and Additional file 1: Figure S5b**). As the chondrocytes originating from the somatic mesoderm are not considered as fibroblasts, we have excluded them from downstream analysis [55]. Fib.origin initially appear at E13.5 and progressively diminish by E15.5, whereas other subclusters begin to appear at E14.5 and subsequently expand. This suggests that Fib.Origin may have the potential to differentiate into various fibroblast subtypes.

To estimate the cellular differentiation dynamics among fibroblast subclusters, we employed RNA velocity, which predicts the future states of individual cells by analyzing the ratios of spliced and unspliced mRNAs [55] (**Fig. 4c**). The RNA velocities indicated a directional flow from the Fib.Origin at the center of the embedding towards the later time points located at the periphery of the embedding (**Fig. 4c**). To further refine our velocity predictions, we applied CellRank, which detects the probabilities of initial and terminal states for each cell based on RNA velocity [56, 57]. Consistent with RNA velocity predictions, CellRank identified higher initial cell states probabilities in Fib.Origin and higher terminal cell states probabilities in other subclusters (**Fig. 4d**).

Next, we further investigated gene expression and corresponding regulatory states along each trajectory. We calculated Pearson correlation between gene expression level and fate probabilities to identify differential genes expressed biased towards Fib.DC fate. Genes with significant positive correlations were defined as candidate driver genes [56]. We identified 527 driver genes in the Fib.DC trajectory, including early markers such as *Fst* and *Sema6a*, as well as late markers such as *Dll1* and *Bmp3* (correlation >0.05 and adjusted *p* value□<□0.05) (**Fig. 4e-g**). To further investigate when and how these driver genes were regulated during fibroblast differentiation, we extracted 10,547 cells alongside the Fib.DC trajectory (fate probability >75% quantile) and sorted by pseudotime ordering. We then performed peak-gene linkage analysis as described above and identified 495 peak-gene pairs (**Fig. 4e**). Using k-means clustering, 527 differential genes were classified into two distinct groups (stage 1 and stage 2, **Fig. 4e**). Genes upregulated at the stage 1 of the trajectory are enriched in pathways such as skin development, epithelial cell proliferation and WNT signaling. These pluripotent functionalities indicate maintaining of Fib.Origin homeostasis. In contrast, genes upregulated at the stage 2 of Fib.DC trajectory are associated with molting cycle, regulation of animal organ morphogenesis and appendage development (**Fig. 4g**). We then applied an analogous approach to the Fib.Inter trajectory and identified 1,196 differential genes, such as *Mfap5* and *Klf4* (**Additional file 1: Figure S5c**). Pathway enrichment analysis of these genes revealed the involvement of mesenchymal cell proliferation in the stage 1, cell aggregation and fibroblast proliferation in the stage 2 and stage 3 of the trajectory (**Additional file 1: Figure S5d, e**)

To examine the *cis*-regulatory landscape of fibroblasts, we defined the underlying GRNs using FigR [31]. We calculated TF motif enrichment, considering both expression level and the chromatin accessibility for all DORCs, to generate the regulation score representing the intersection of motif-enriched and RNA-correlated TFs. We distinguished five unique modules of DORCs that are regulated by TFs (**Additional file 1: Figure S5f**). key TFs for fib.Origin are enriched in module 2 subclusters, such as WNT/β-catenin associate factor LEF1 [58, 59] (**Additional file 1: Figure S5g).**

To further identify potential regulatory gene targets of TFs driving Fib.DC cell identify, we extracted the cells from the Fib.DC trajectory and examined gene regulatory networks using FigR. We calculated the Spearman correlation between gene expression and accessibility of peaks within a 100□kb window around TSS (*p* value < 0.05). Then we identified 395 DORC regions (each with at least four significant peak-gene associations) and 125 TF motifs (regulation score ≥ 1), and determined two distinct gene modules regulated by different TFs (**Fig. 4h and Additional file 1: Figure S5h**). We proposed several potential regulatory factors that could serve as lineage determinants for Fib.DC and DP, including RUNX2, SOX2, ALX4 and PRRX2 (**Fig. 4h and Additional file 7: Table S7**). ALX4 plays a crucial role in hair follicle development, as *Alx4*-null mice exhibit dorsal alopecia [60]. SOX2 as a key TF regulates hair growth by modulating WNT signaling [61]. Next, we applied CellOracle to simulate changes in Fib.DC identity upon TF perturbation [62]. This silico strategy employs GRN to simulated cell state of each cell following perturbation of candidate TFs. We then calculated perturbation scores for TFs detected in FigR (**Fig. 4i and S Additional file 8: Table S8**). High perturbation scores suggest that in silico knockout of the TF significantly decreased, suggesting that the TF is an essential regulator of Fib.DC trajectory. Interestingly, while many TFs showed correlated perturbation scores according to CellOracle, RUNX2, ALX4 and PRRX2 exhibited relatively high specificity for Fib.DC trajectory (**Fig. 4j**).

## Discussion

Here we present a single cell multimodal profiling approach named HT-scCAT-seq, which enables joint detection of transcriptome and chromatin accessibility for ten thousand of single-cells in one time. We benchmark HT-scCAT-seq data with other single-cell joint-profiling approaches such as 10x Genomics and ISSAC-seq and find that HT-scCAT-seq produced data with equal quality or even better quality comparing with similar approaches. Finally, we applied HT-scCAT-seq to mouse embryonic skin samples, to dissect the underlying transcriptional regulatory program for each cell lineage. By TF-peak-gene linkage analysis, we demonstrated how lineage specific chromatin regulators facilitate downstream gene expression and reconstructed a lineage specific chromatin regulatory network for Fib. Origin subtype.

HT-scCAT-seq is a stable and easy handling single-cell method for ordinary biological and medical laboratories. Although experiments in this work are performed with DNBelab C4 platform, HT-scCAT-seq can simply adapt to other microfluidic based devices such as 10x Chromium, Bio-Rad ddSEQ or manual ones. The throughput of HT-scCAT-seq is largely between one to ten thousand per reaction, and can be scale up to a hundred thousand by employing combinatory-indexing strategy which introduce an extra cellular barcode in transposition and reverse-transcription steps. Besides, these extra barcodes can be served as sample barcodes, which may be time-saving, cost-efficient and potentially free from batch effect.

Recent studies have shed light on the idea that chromatin structure is a key determinant for gene expression [63–66]. Factors such as binding of transcription factors, chromatin accessibility, nucleosome occupancy, DNA methylation and histone modifications play roles in building chromatin landscape. It is imprecise to link two populations defined by different omics data, therefore multi-omics data can give us direct linkage of two or more layers of the regulome [67]. Yet current experimental methods including HT-scCAT-seq are still limited in profiling two or three layers in one time, which is still a big challenge for researchers. However, in-silico assembly of multiply datasets containing different layers together may help us to reconstruct the panorama of epigenetic state, which may revolute our understanding of cell types and cell states.

## Conclusions

In this study, we introduce an enhanced single-cell multimodal profiling technique, HT-scCAT-seq, which simultaneously detects chromatin accessibility and gene expression in individual nuclei in high throughput manner. We highlight the efficacy and cost-efficiency of HT-scCAT-seq as a high-throughput method for single-cell multiomics. We provide extensive evidence of cell-type-specific candidate *cis*-regulatory elements (cCREs) and transcription factors (TFs) that are essential for decoding the transcriptional regulatory program of mouse embryonic skin.

## Methods

### Cell culture

HEK293T and NIH3T3 were cultured in DMEM, high glucose (Thermo, 11965126) supplemented with 10% FBS (HyClone™, SH30071.03) and 1% Penicillin-Streptomycin (Thermo, 10378016) at 37°C with 5% CO_2_.

### Animal study

Wild-type C57BL/6 mice (Guangdong medical laboratory animal center) were interbred and pregnant females were sacrificed at E13.5, E14.5, E15.5, E16.5 and E18.5. Embryonic dorsolateral skin was micro-dissected, pooled (5 embryos per time point) and pre-chilled in cold PBS, followed by cell dissociation and nuclei extraction steps. Total twenty-five mouse embryos were processed and analyzed in following experiments. Mouse brain tissue was collected from wild-type C57BL/6 mice aged 8 weeks.

### Cell dissociation from embryonic dorsolateral skin

To generate a single cell suspension, embryonic dorsolateral skin samples were first digested by Enzyme I (1% Penicillin-Streptomycin (Thermo, 10378016) and 0.125% Trypin (Thermo, 25200056)), which were incubated at 37°C for 10 min on a thermo shaker. A mixture of tissue and cells was filtered using a 70 µm cell strainer (Falcon, 352350). The obtained mixture was incubated Enzyme II (2.5 mg/mL Collagenase Type IV (Thermo, 17104019), 1 mg/mL Collagenase Type I (Thermo, 17100017), 1 mg/mL DispaseII (Thermo, 17105041), 1% Penicillin-Streptomycin (Thermo, 10378016)) in DMEM/F-12 (Thermo, 11320033) for 15 min at 37°C on a thermo shaker. Dissociated cells were filtered through a 40 µm cell strainer (Falcon, 352340). Cell suspension was centrifuged for 5 min at 500 g and cells were washed with 0.04% BSA/PBS for 1 or 2 times.

### Nuclei preparation and fixation

For the species mixing and embryonic dorsolateral skin experiments, single-nucleus preparations were derived from the Omni-ATAC protocols as previously described [68], with some adjustments. In brief, 5 x 10^5^ cells were collected and resuspended in 100 µL of chilled cell lysis buffer (10 mM Tris-HCl pH7.5, 10 mM NaCl, 3 mM MgCl_2_, 0.1% Tween-20 (Sigma, P9416), 0.1% NP40 (Roche, 11332473001), 0.01% digitonin (Sigma, D141), 1% BSA/PBS and 0.8 U/μL RNase inhibitor (Neoprimaries, LS-EZ-E-00006P), and incubated on ice for 5 min. Subsequently, 1 mL of chilled resuspension buffer (10 mM Tris-HCl pH7.5, 10 mM NaCl, 3 mM MgCl_2_, 0.1% Tween-20, 1% BSA/PBS and 0.8 U/μL RNase inhibitor) was added into the lysed cell suspension, and nuclei were spun down at 500 g, 4°C for 5 min. Nuclei were resuspended in 50 µL of PBSI (1 U/μL RNase inhibitor and 0.5 U/μL SUPERase inhibitor (Thermo, AM2696)) and counted using DAPI staining.

For the adult mouse brain experiments, single-nucleus preparations were derived from the Omni-ATAC protocols as previously described [68]. Mouse brain tissue was placed into a pre-chilled 2 mL Dounce homogenizer with 2 mL of homogenization buffer (HB; 1 x basic buffer (20 mM Tris pH 7.8, 25 mM KCl and 5 mM MgCl_2_) 250 mM sucrose, 1 mM DTT, 1 x Protease Inhibitor Cocktail, 0.8 U/μL RNase inhibitor and 1% BSA). Tissue was homogenized with 15 strokes using pestle A and the homogenate was filtered through a 70 μm cell strainer. Transfer homogenate to a new Dounce homogenizer and homogenize with 10 strokes using the pestle B. The homogenate was filtered through a 40 μm cell strainer and centrifuged at 500 g, 4°C for 5 min. 2.4 mL of nuclei wash buffer (20% iodixanol, 1 x basic buffer in HB) were added into the nuclei suspension, and nuclei were spun down at 800 g, 4°C for 10 min. Nuclei were resuspended in 50 µL of PBSI and counted by DAPI staining.

Nuclei were fixed in 0.1% formaldehyde (Sigma, 1209228) in PBSI for 5 min at room temperature and quenched with 0.125 M glycine (Sigma, 635782). Fixed nuclei were spun down at 500 g, 4°C for 5 min, followed by washing twice in PBSI and resuspended in 50 µL of PBSI.

### HT-scCAT-seq library preparation and sequencing

For in situ transposition, 200,000 fixed nuclei were resuspended in tagmentation mix (0.08 U/μL Tn5 transposase and 1 x TAG buffer in PBSI) and incubated at 37°C for 30 min with shaking (500 rpm). The nuclei were then centrifuged at 500 g, 4°C for 5 min and resuspended in 8 µL PBSI. Transposed nuclei were mixed with reverse transcription buffer (0.625 mg/mL TransFlex III Reverse Transcriptase (Neoprimaries, LS-EZ-E-00027Q), 1 x RT buffer, 1 mM dNTP mix, 2.5% PEG 6000, 1 mM dCTP, 3.75 μM Oligo dT, 2.5 μM TSO and 1 U/μL RNase inhibitor), and RT was performed (10°C for 30 s, 20°C for 30 s, 30°C for 30 s, 40°C for 30 s, 50°C for 5 min, 4°C for 3 min, 10°C for 45 s, 20°C for 45 s, 30°C for 30 s, 42°C for 2 min, 50°C for 10 min). After in situ reverse transcription, nuclei were centrifuged at 500 g, 4°C for 5 min, washed twice, and resuspended in 50 µL of 1% BSA in PBS. Libraries were generated using DNBelab C Series Single-Cell ATAC Library Prep Set (MGI, 940-000793-00) following the user protocol with the following modifications: in encapsulation and pre-amplification step, the RNA PCR primer (biotin-modified) was added; after emulsion breakage, MyOne C1 Dynabeads were added to an equal volume mixture of RNA and ATAC product with 1 x B&W buffer (50 mM Tris pH 7.5, 0.5 mM EDTA, 1M NaCl); after emulsion breakage, RNA and ATAC product were purified and constructed separately. All libraries were sequenced using DIPSEQ T1 platform at China National GeneBank (CNGB).

### Single-cell data preprocessing

In the initial preparation of single-cell ATAC-seq (scATAC-seq) data, we first aligned Read 1 and Read 2 FastQ files to either the mm10 or hg38 genome. We then used Chromap (v0.2.1) [22] to generate fragment files. To process the barcodes, we utilized d2c (v1.5.3) to calculate and merge beads from the same barcode. MACS2 (v2.28) [69] was employed to identify peaks and produce a peak matrix. For the scRNA-seq data, Read 1 FastQ (comprising Barcode 1, Barcode 2, and UMI) and Read 2 FastQ were aligned and annotated using scStar (v1.0.3) and Anno (v1.4). Beads were subsequently merged into barcodes based on the results from the d2c step in the ATAC-seq data preprocessing, and the gene expression matrix was derived using PISA (v1.10.2) [24].

For the species mixing experiment, we combined HEK293T and NIH3T3 cells and processed them through the HT-scCAT-seq workflow utilizing a combined reference genome of hg38 and mm10. Cell barcodes with more than 80% of reads mapped exclusively to a single genome were classified as singlets; others were considered as doublets.

### Single-cell metric comparison to other methods

We downloaded FastQ files of cell line data from other multi-omics technologies. To ensure uniformity, we downsampled all data sets to 50,000 read pairs per cell for each modality. We then applied the ISSAAC preprocessing workflow to these data consistently. For scATAC-seq data, we employed Chromap (v0.2.1) [22] for read mapping. Meanwhile, scRNA-seq reads were analyzed using STARsolo (v2.7.10a) [70]. From these processed data, we derived count matrices which were subsequently utilized by ArchR (v1.0.2) [71] to calculate quality metrics.

### Mouse brain data analysis

The gene expression matrix of mouse brain was processed using Seurat (v4.3.0.1) [36] to create a Seurat object. Cells were filtered based on the following metrics: nCount_RNA between 200 and 100,000, nFeature_RNA less than 7,500, and percent.mt less than 20. Each data modality was then adjusted for batch effects using Harmony. Data normalization was performed using the NormalizeData function. To identify top 3,000 variable genes, the FindVariableFeatures function was used. The first 50 principal components (PCs) were determined by running RunPCA on these variable genes, and cell clusters were identified using FindNeighbors. The FindAllMarkers function was used to identify marker genes in each cluster. For mouse brain scATAC-seq data, the peak matrix was processed using Signac package (v1.12.0). Cells were filtered based on the following metrics: nCount_ATAC between 2,000 and 20,000, FRiP greater than 0.2, and TSS.enrichment between 2 and 20. The matrix was normalized, and clusters were identified with dimensions ranging from 2 to 40, using default settings according to the Signac documentation. The difference of the chromatin status between clusters were computed by peak calling using MACS2.

### Prediction of RNA expression by DORC, SCARlink and Signac

Paired cells of scATAC-seq and scRNA-seq mouse brain data was used in these assays. High density DORCs were calculated within a distance of 50 kb from each gene’s TSS, and the spearman correlation coefficient was calculated for each gene-peak pair [31]. ChromVAR (v1.16.0) [44] was used to generate the overall accessibility and GC content, which was then used to perform background peak correction. One-tailed *Z*-test was calculated to determine the association of each gene-peak pair. DORCs were then defined as those with a permutation *p* value less than or equal to 0.05 and genes with more than four associated peaks in this assay. To obtain a single-cell DORC score matrix, the scATAC-seq data was first normalized by peak counts. Each gene’s DORC score was then calculated as the sum of counts based on the significantly correlated peaks per gene. SCARlink [33] was applied to the 250 kb upstream and downstream of the gene body. Regularized poisson regression was performed to predict gene expression. Top 3,000 variable genes were selected using Seurat [36], which were used as input to SCARlink. Predicted gene expression matrix was generated using SCARlink with default parameters.

### Mouse embryonic skin data analysis

The processing of single-cell RNA and ATAC data of mouse embryonic skin was using Seurat [36] and Signac [32]. Quality control is performed separately for scRNA-seq and scATAC-seq data. Cells with high mitochondrial content (percent.mt > 20) or low RNA counts (nCount_RNA < 200) are removed from the scRNA-seq dataset. For scATAC-seq, cells with TSS enrichment score less than 4 and fewer than 2,000 captured fragments are excluded.

In both scRNA and scATAC, we used scDblFinder (v1.16.0) [72] and Harmony to remove doublet and batch effect. For scRNA data, we also performed removal of ambient RNA by using SoupX (v1.6.2) [73] with default parameters. after background filtration, normalization and scaling of RNA gene expression levels was performed using the NormalizeData and SCTransform function. Then, PCA is performed, and the first 50 principal components were used to group cells into clusters using FindNeighbors and FindClusters functions in Seurat. For scATAC data, iterative LSI dimensionality reduction is performed for 2:40 components, taking the top 50% variable peaks and evaluating resolutions 0.5. Cells are then projected in a 2D space using the RunUMAP function. The scATAC-seq data was annotated using the corresponding annotation of paired scRNA-seq cells. Gene activity matrix was created with function GeneActivity in Siganc. Motif activity matrix was calculated using chromVAR based on JASPAR 2020 database, and FindMotifs function was used to identify differentially enriched motifs, TF motifs with interest were visualized and used for further analysis [74].

### Peak-to-gene linkage identification

Same as the description in the ‘prediction of RNA expression by DORC’. Briefly, the associations between peak and gene were calculated using LinkPeaks function in Signac. The Pearson correlation coefficient was calculated for each peak within a distance of 50 kb from the gene’s TSS. *p* value was then adjusted by Benjamini-Hochberg method. Peak-gene links with coefficients < 0 and adjusted *p* value > 0.05 were removed [31]. This ensures that only statistically significant and positively correlated peak-gene pairs are considered for downstream analyses.

### Identification of fibroblast subtypes

We extracted all fibroblast cluster cells from mouse embryonic skin data and utilized R package Seurat and Signac to conduct a general upstream analysis. Briefly, 2,000 variable genes and a maximum dimension of 40 were selected. The SLM algorithm and a resolution of 0.5 was used in FindClusters functions to produce cell-type clusters, which we then annotated based on marker gene expression. To identify DEGs and differential peaks, Seurat FindAllMarkers with default setting was performed. We used clusterProfiler (v3.11.0) to enrich Gene ontology (GO) terms for differentially expressed genes [75]. Additionally, top 500 stage-specific peaks were visualized by heatmap in periderm cells, and were imported into the GREAT analysis website (http://great.stanford.edu/public/html/) for GO enrichment analysis [49].

### RNA velocity and trajectory inference

Data of two modalities processed by Seurat and Signac were loaded as “AnnData” object with Chod cell type removed. Spliced and unspliced reads information was extracted from possorted bam file using velocyto (v0.17.15) and added to the object [76]. The data was then filtered and normalized using scVelo.pp.filter_and_normalize function with parameters (min_shared_counts=20, n_top_genes=3000), 40 principal components were used to find the cell neighbors using scVelo.pp.neighbors. Velocity analysis was performed using ‘scVelo.tl.velocity’ with default parameters. Previous UMAP coordinates was used for velocity visualization.

We used scanpy.tl.diffmap to designate a random cell of Fib.Origin as the root cell and calculated pseudotime using scanpy.external.palantir. Subsequently, we computed a directed cell-cell transition matrix using the PseudotimeKernel from Cellrank (v2.0.4) [77]. We then calculated and visualized fate probabilities towards terminal states based on the GPCCA module established by PseudotimeKernel. We used g.compute_lineage_drivers to compute driver genes, and filtered driver genes with correlation greater than 0.05 and adjusted *p* value less than 0.05. These driver genes were calculated for associated peaks with LinkPeaks in Siganc, and the result was visualized by heatmap. For the Fib.DC lineage, we set the first 95% of genes as start/middle and the last 5% of genes as end for GO enrichment analysis.

### TF-DORCs regulatory network

DORCs were determined as the above description (prediction of RNA expression by DORC), DORC score matrix was obtained and smoothed using cisTopic (v0.3.1) with the LSI algorithm. TF-DORCs associations was calculated using runFigRGRN in FigR (v1.0.1) with regulation score ≥ 2 retained [31]. The final networks were constructed using igraph, with node and edge attributions formed according to gene expression and chromatin accessibility.

### CellOracle analysis

To validate the identified TFs, we performed the in silico perturbation analysis using CellOracle (v0.18.0) [62]. First, we utilized Cicero (v1.20.0) to identify distal cis-regulatory elements [78]. Subsequently, we annotated the co-accessible peaks and filtered out active promoter/enhancer elements. Next, we constructed a cell type-specific GRN for the TF scan, retaining network edges with *p* value above 0.001 and top 2000 edges. Finally, we set the TF expression as 0 to compute the perturbation scores in different clusters.

## Declarations

### Ethics approval and consent to participate

All mice experiments were approved by the Institutional Review Board on the Ethics Committee of BGI.

### Consent for publication

Not applicable.

### Availability of data and materials

All raw data have been stored in CNGB Nucleotide Sequence Archive (CNSA) [79] of China National GeneBank DataBase (CNGBdb) [80], with accession number CNP0005787. All data were analyzed using standard programs and packages, as detailed above. Source code and analysis scripts supporting the findings of this study are available on the Github repository (https://github.com/caipf/HT-scCAT-seq).

### Author contributions

C.L. and L.L. conceived the idea; C.L. and Y.Y. supervised the study; Y.Y., Q.D. and Y.L. designed the experiment; Q.D. and Y.Y. performed the majority of the experiments with the help of Z.H., J.X., M.C., X.L., R.Z., S.D., J.C., R.L., X.S. and Chang L.; P.C. and Z.Z. analyzed the data with the help of W.M., X.C., S.H. and W.M; P.G., J.L. and Jun X. provided project support; Q.D., Y.L. and P.C. wrote the manuscript; C.L. and Y.Y. participated in the manuscript editing and discussion.

## Acknowledgements

We thank all our teams’ members and the China National GeneBank (CNGB) for their support.

## Additional files

**Additional file 1:** Figures S1-S5

**Additional file 2:** Tables S1-S2

**Additional file 3:** Table S3

**Additional file 4:** Table S4

**Additional file 5:** Table S5

**Additional file 6:** Table S6

**Additional file 7:** Table S7

**Additional file 8:** Table S8

## Additional file 1: Fig S1-S5

**Fig S1.**
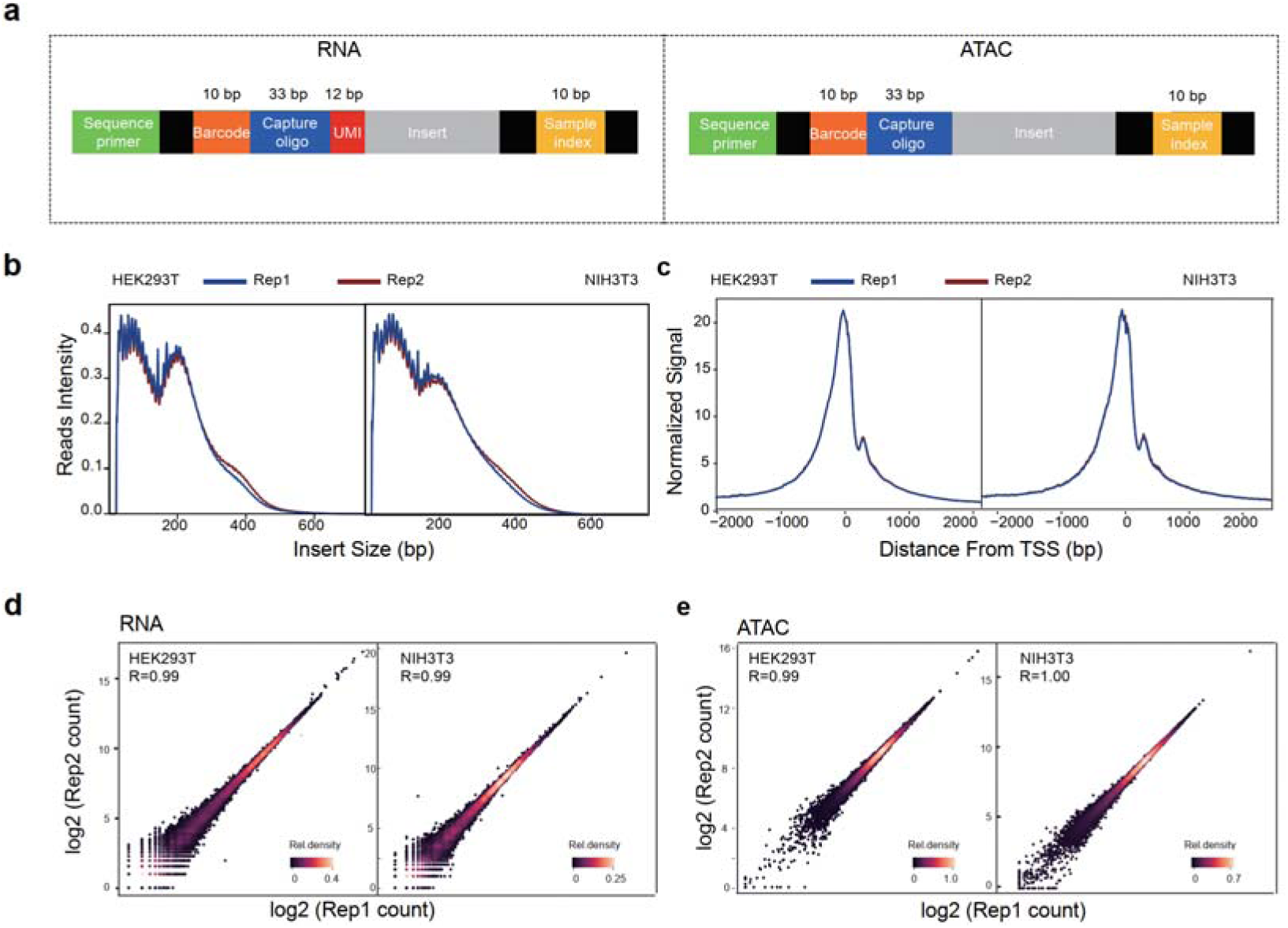
Quality control for HT-scCAT-seq data on the cell line. a The structure of the HT-scCAT-seq RNA and ATAC-partition library. **b-c** The insert size distribution (b) and the enrichment read around TSSs (c) in ATAC data from HEK293T cells (left) and NIH 3T3 cells (right). **d-e** Scatter plots showing the reproducibility of ATAC data (d) and RNA data (e) from two biological replicates.

**Fig S2.**
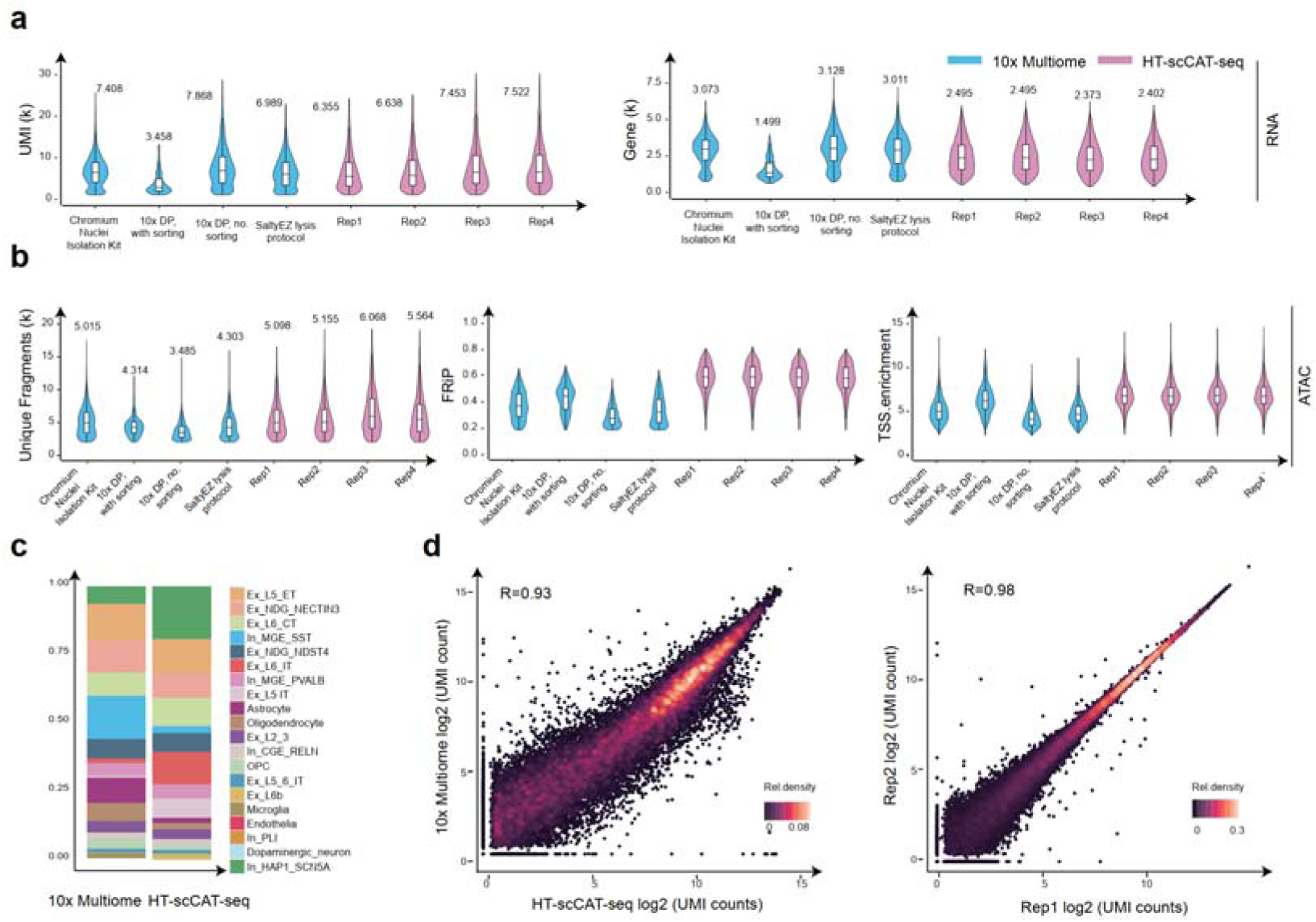
Quality control for HT-scCAT-seq data on the mouse brain. **a-b** Violin plots showing the distribution of UMIs (a, left panel), detected gene number (a, right panel), fragment (b, left panel), FRiP (b, middle panel) and TSS enrichment (b, right panel) between 10x Multiome and HT-scCAT-seq platforms. **c** Fraction of cells in each cell type from RNA dataset of two platform (10x Multiome and HT-scCAT-seq). **d** Scatter plots showing the reproducibility of RNA data from two platform (left panel, 10x Multiome and HT-scCAT-seq) and two biological replicates (right panel, HT-scCAT-seq).

**Fig S3.**
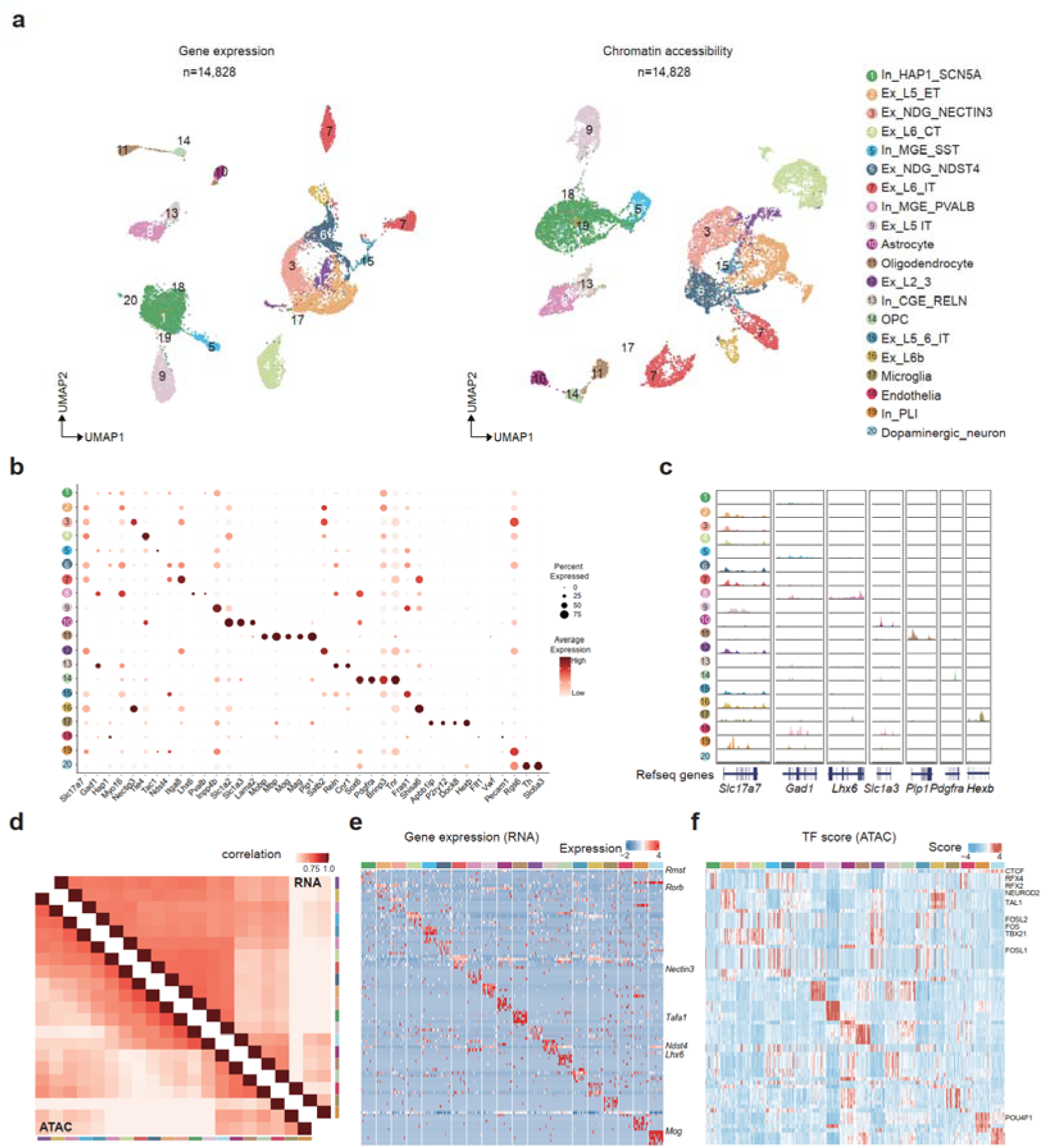
Data quality for HT-scCAT-seq data on the mouse brain. **a** UMAP visualization of RNA (left) and ATAC (right) profile from 14,828 mouse brain nuclei. **b-c** Bubble plot showing gene expression level (b) and aggregate track showing ATAC signal (c) of selected genes. **d** Spearman correlation coefficient between cell types in RNA (top right) and ATAC (bottom left) datasets. For RNA data, the correlation was calculated using the average gene expression per cell type, while the ATAC data correlation was calculated based on the average peak counts per cell type. **e-f** DEGs expression (e) and differential TF motif scores (f) of each cell type.

**Fig S4.**
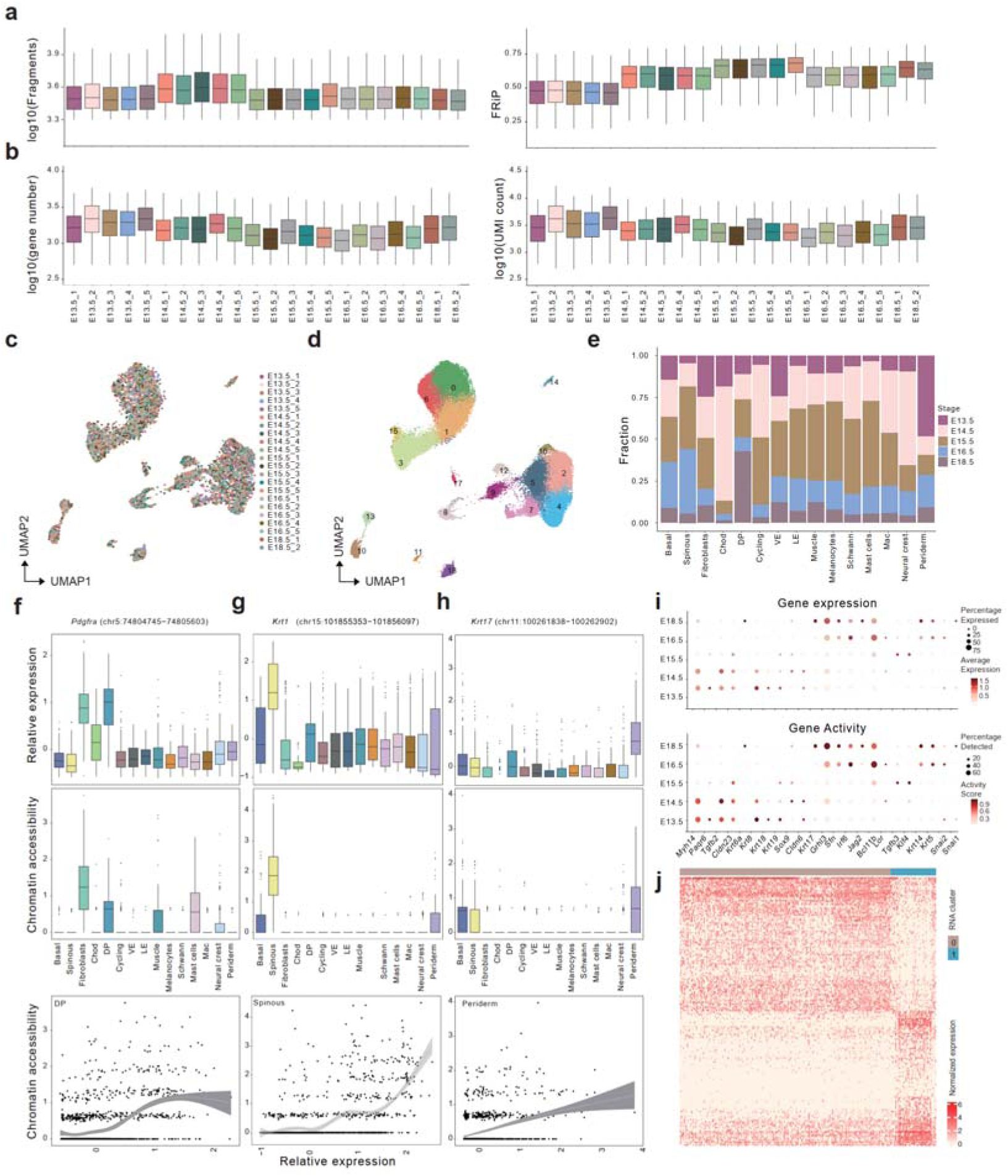
HT-scCAT-seq generates high quality on mouse embryonic skin, related to Figure 3. **a-b** Box plots showing distribution of several ATAC (a) and RNA (b) metrics for each library. ATAC metrics include number of unique fragments (left) and FRiP (right). RNA metrics include number of detected gene (left) and UMIs (right). **c-d** UMAP projection of full RNA dataset, colored by experimental samples (c) and clusters (d). **e** Bar graph displaying cell type composition of each developmental stages. **f-h** Boxplots showing expression levels (top) and relative accessibility (bottom) of Pdgfra (f), Krt10 (g) and Krt17 (h) across different cell types. **i** Bubble plot showing gene expression level (top) and gene activity scores (bottom) of selected genes in periderm cell. **j** Heatmap showing gene expression of top 2,000 cluster-specific genes of periderm subtypes.

**Fig S5.**
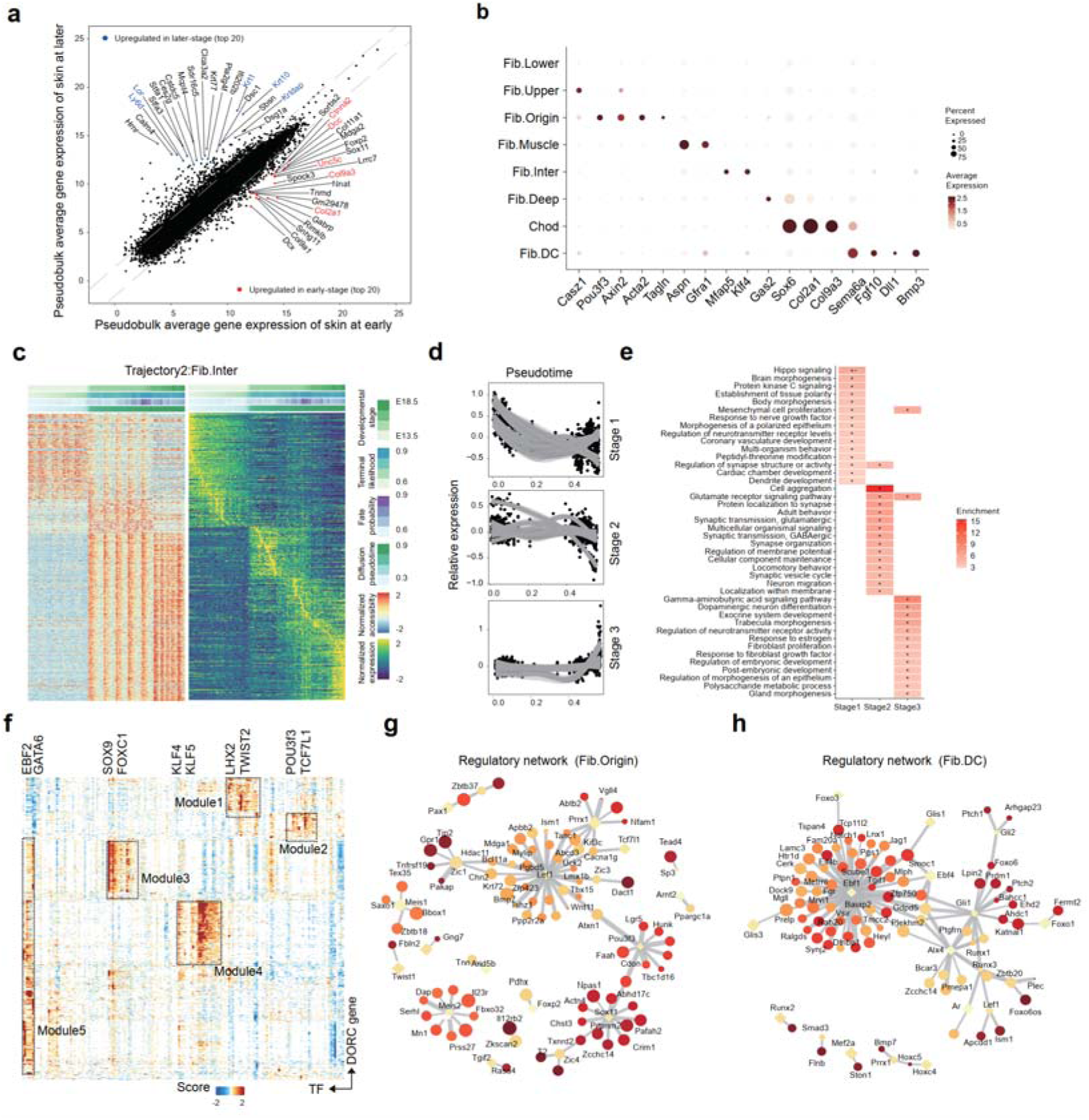
Fibroblast trajectory analysis and TF regulatory network construction. **a** scatter plots show DEGs in the fibroblasts in early (E13.5 – E14.5) and late stage (E15.5 – E18.5). Top 20 genes are labeled. **b** Dot plot showing gene expression values of selected genes in fibroblasts data. **c** Heatmap showing chromatin accessibility (left) and gene expression (right) of 1,767 linked peak-gene pairs across 9,138 cells in the Fib.Inter trajectory. Each row represents a linked gene and a pair of cCREs. Bars on the top represent diffusion pseudotime, fate probabilities to the anterior palatal mesenchymal trajectory, terminal state likelihood, and developmental stage. Columns are ordered by diffusion pseudotime. **d** scatter plots show the correlation between gene expression values (y-axis) and pseudotime (x-axis) along the Fib.Inter trajectory. The fitted smooth line indicates a 95% confidence interval. **e** Top 15 enriched Go terms for drive genes upregulated at the start, middle and end of the Fib.lnter trajectory are presented. Dot size denotes the enrichment ratio and color reflects the level of significance. **f** Heatmap shows TF-DORC regulation scores derived from fibroblast dataset using FigR (n = 161 TFs, n = 900 DORCs). Each row represents a DORC, while each columns represents a TF. **g** Visualization of the TF-DORC network formed by Fib.Origin module (module 2). DORCs are depicted as circles, and associated TFs as diamonds. The size of circles and the intensity of colors indicate the level of DORC expression. **h** Same as in **g,** but for Fib.DC module (module 2 in Fig. 4h).

